# Chromatin regulator *Kdm6b* is required for the establishment and maintenance of neural stem cells in mouse hippocampus

**DOI:** 10.1101/2024.02.20.581302

**Authors:** Eugene Gil, Sung Jun Hong, David Wu, Dae Hwi Park, Ryan N. Delgado, Martina Malatesta, Sajad Hamid Ahanger, Karin Lin, Saul Villeda, Daniel A. Lim

**Author notes:** equal contribution (Eugene Gil, Sung Jun Hong).

## Abstract

Neural stem cells (NSCs) in the mouse hippocampal dentate gyrus (DG) – a structure important to learning and memory – generate new neurons postnatally and throughout adult life. However, the regulators that enable this lifelong neurogenesis remain incompletely understood. Here we show that the chromatin regulator KDM6B is required for both the establishment and maintenance of NSCs in the mouse DG. Conditional deletion of *Kdm6b* in embryonic DG precursors results in an adult hippocampus that is essentially devoid of NSCs, and hippocampal-dependent behaviors are defective. *Kdm6b*-deletion causes precocious neuronal differentiation, and the NSC population fails to become established in the postnatal DG. Using single cell RNA sequencing (scRNA-seq), we observed that *Kdm6b*-deletion disrupts the transcriptomic signature of NSC maintenance. Furthermore, deleting *Kdm6b* in adult DG NSCs induces early neuronal differentiation, and the NSC population is not properly maintained. These data illustrate the critical role that *Kdm6b* plays in adult DG neurogenesis, which may help understand how mutations in this chromatin regulator result in cognitive disorders in human patients.

## INTRODUCTION

In most mammals, a local population of neural stem cells (NSCs) is established and maintained in select regions of the adult brain, enabling neurogenesis to persist throughout life (Kempermann et al., 2015; Ming and Song, 2011). In mice, NSCs become established in the dentate gyrus (DG) in early postnatal development and are maintained throughout adulthood (Berg et al., 2019). Although the transcriptomic signature of mouse DG NSC maintenance has been described (Shin et al., 2015; Hochgerner et al., 2018), and several transcription factors have been identified as being involved in the long-term function of DG NSCs (Ables et al., 2010; Imayoshi et al., 2010; Engler et al., 2019; Lampada and Taylor, 2023), less is known about the chromatin regulators that orchestrate the transcriptome required for the establishment and maintenance of NSCs in the adult hippocampus.

Lysine (K) Demethylase 6B (KDM6B, also known as Jumonji domain-containing protein D3 (JMJD3)), is a chromatin regulator vital for proper neural development (Burchfield et al., 2015). Mutations in *KDM6B* have been linked to deficits in cognitive function, and *KDM6B* has been identified as a as a risk gene for autism spectrum disorder (ASD) (De Rubeis et al., 2014; Iossifov et al., 2014; Sanders et al., 2015; Satterstrom et al., 2020; Rots et al., 2023). In addition to its role as a demethylase specific for histone 3 lysine 27 trimethylation (H3K27me3) (Agger et al., 2007; Burchfield et al., 2015; Hong et al., 2007) – a chromatin modification associated with transcriptional repression (Di Croce and Helin, 2013; Margueron and Reinberg, 2011) – KDM6B can also regulate transcription via demethylase-independent activities (Miller et al., 2010; Vicioso-Mantis et al., 2022). In mouse embryonic stem cells, *Kdm6b* is required for neural induction (Burgold et al., 2008), and expression of *Kdm6b* is regulated during forebrain development (Jepsen et al., 2007). Knockdown studies of *Kdm6b* in the chicken embryonic spinal cord and mouse retina further demonstrate critical roles for this chromatin regulator in neurogenesis of the developing central nervous system (CNS) (Akizu et al., 2010; Iida et al., 2014). *Kdm6b* has also been shown to be important for induction of genes linked to inflammation in synaptically activated hippocampal neurons (Wijayatunge et al., 2014).

In the adult mouse ventricular-subventricular zone (V-SVZ) – another germinal zone that produces large numbers of new neurons throughout life – *Kdm6b*-deleted V-SVZ NSCs are defective for neuronal differentiation, failing to properly activate key neurogenic transcription factors (Park et al., 2014). However, the initial establishment of V-SVZ NSCs is not compromised by deletion of *Kdm6b* (Park et al., 2014).

In this study, we found that *Kdm6b* is required for the establishment and maintenance of NSCs in the mouse DG. Without *Kdm6b*, DG NSCs undergo precocious neuronal differentiation, depleting the NSC population, which produces an adult DG essentially devoid of neurogenesis. Instead of being required to activate genes for neuronal differentiation, *Kdm6b* in DG NSCs is required for the expression of a transcriptomic signature of stem cell maintenance. The perturbation in the transcriptome that results from the deletion of *Kdm6b* appears to drive stem cells into a more differentiated state. Together, these results highlight the distinct roles that *Kdm6b* plays in different NSC populations in the mouse brain and adds to our developmental understanding of the cognitive deficits observed in patients with *KDM6B* mutations.

## RESULTS

### *Kdm6b* is expressed during mouse postnatal DG development

In the subgranular zone (SGZ) of the adult mouse hippocampal DG (**Figure 1A**), NSCs produce intermediate-progenitor cells (IPCs), which give rise to neuroblasts that migrate into the granule cell layer (GCL) and become excitatory granule neurons (**Figure 1B**). The SGZ NSC population arises from embryonic radial glial cells – the NSCs of the developing brain – located in a region of the ventricular zone called the dentate neuroepithelium (DNE, **Figure 1—figure supplement 1A**) (Rolando and Taylor, 2014). *In situ* hybridization (ISH) revealed *Kdm6b* expression in the embryonic day 16.5 (E16.5) DNE and developing DG of postnatal day 1 (P1) and P7 (**Figure 1—figure supplement 1B-D**). By the end of the first postnatal week, NSCs localize to the inner layer of the DG (Nicola et al., 2015), and *Kdm6b* was detected throughout the DG at postnatal day 1 (P1) and 7 (P7) (**Figure 1—figure supplement 1C, D**). As postnatal development continues, these NSCs consolidate into the SGZ, and *Kdm6b* expression continued to be observed at P21 (**Figure 1—figure supplement 1E)**. Nuclear KDM6B was also detected by immunohistochemistry (IHC) in essentially all DG cells including the SGZ NSCs (**Figure 1C**). Thus, *Kdm6b* is expressed in the developing DG when SGZ NSCs are being established.

**Figure 1.**
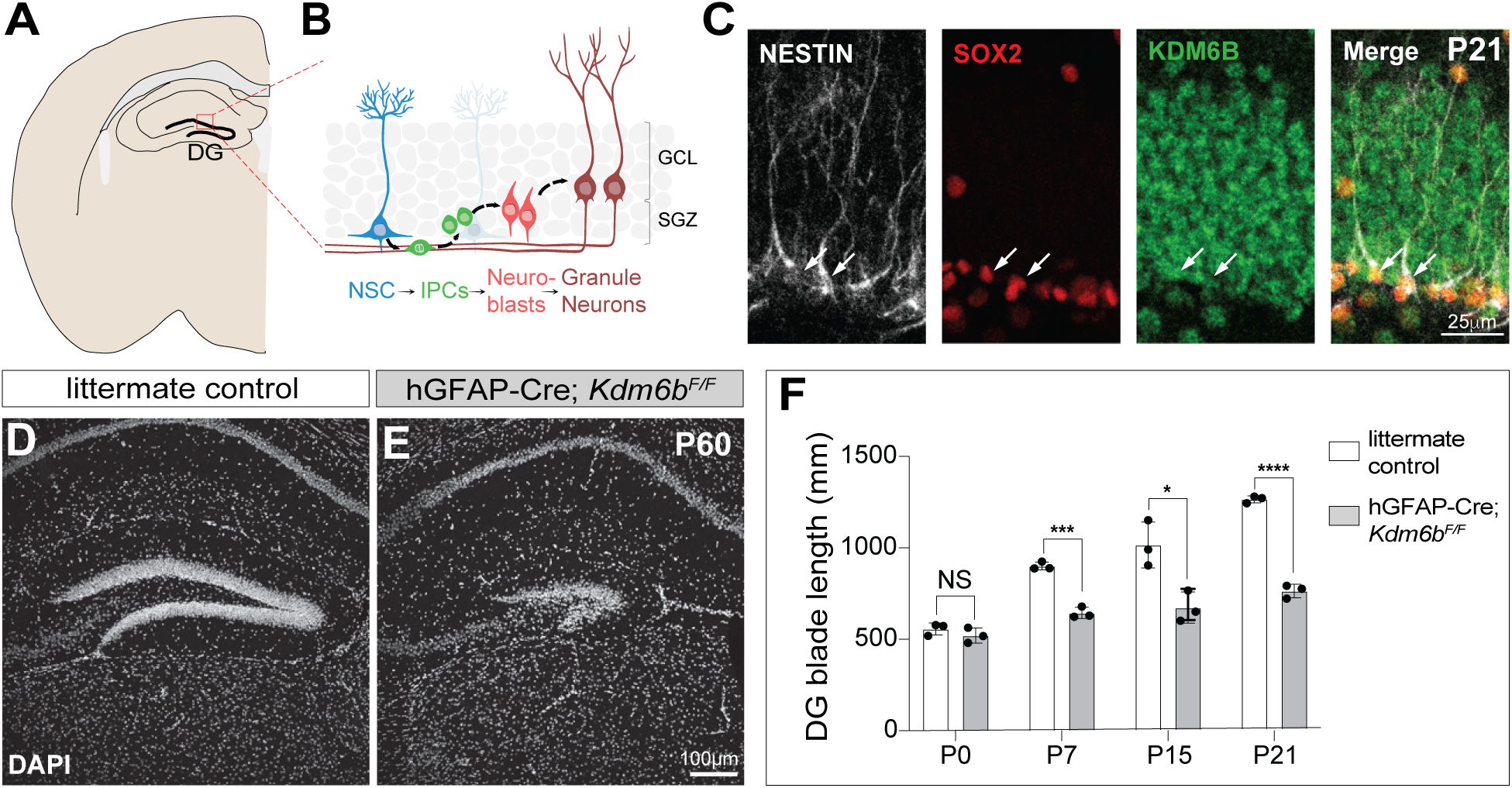
Postnatal DG development is impaired with *Kdm6b*-deletion. **A,** Schematic coronal section showing the dentate gyrus of the hippocampus; red box indicates regions shown in (B). **B,** Schematic illustration of neurogenesis in the DG. **C,** KDM6B (green) expression in the DG of P21 mouse brains. Immunohistochemistry (IHC) is shown for SOX2 and NESTIN. **D-E**, IHC for DAPI (white) in coronal DG sections of P60 control (D) and *hGFAP-Cre;Kdm6b^F/F^* mice (E). **F,** Quantification of length of the DG in control and *hGFAP-Cre;Kdm6b^F/F^* mice from P0 and P21 (n = 3 each), two-tailed unpaired t test (* = p < 0.05, *** = p < 0.0005, **** = p < 0.0001, NS = not significant).

### *Kdm6b* is required for postnatal DG development

To investigate the role of *Kdm6b* in DG development, we targeted *Kdm6b-*deletion to the developing hippocampus by crossing mice expressing Cre under control of the human glial fibrillary acidic protein promoter (*hGFAP-Cre*) with mice carrying conditional knockout alleles of *Kdm6b* (*Kdm6b^F/F^*) (Iwamori et al., 2013; Park et al., 2014). *hGFAP-Cre* is expressed in the hippocampal ventricular zone and results in efficient recombination in cells of the DNE and developing DG by E16.5 (Han et al., 2008). As expected, the DNE and DG of *hGFAP-Cre;Kdm6b^F/F^* mice lacked wild-type *Kdm6b* transcripts as detected by ISH (**Figure 1—figure supplement 1F, G**). *hGFAP-Cre;Kdm6b^F/F^*mice and their littermate controls (*Kdm6b^F/+^, Kdm6b^F/F^* and *hGFAP-Cre;Kdm6b^F/+^*) were born at the expected Mendelian ratios and were similar in overall size, weight and survival throughout postnatal and adult life (Park et al., 2014).

The DG continues to grow after birth, and by P21, it reaches a size similar to that of adult mice (Nicola et al., 2015). At P0, the developing DG in *hGFAP-Cre;Kdm6b^F/F^*mice was similar to littermate control mice. While the DG in control mice exhibited progressive increase in its blade length from P0 to P21, the DG in *hGFAP-Cre;Kdm6b^F/F^*mice did not grow at the same rate, being significantly smaller at every time point examined from P7 to P21 (**Figure 1F**). In adult (P60) *hGFAP-Cre;Kdm6b^F/F^* mice, the DG was very small and hypocellular as compared to littermate controls (**Figure 1D, E)**. Thus, without *Kdm6b*, postnatal mouse DG development is impaired.

### Hippocampal-dependent behavior is defective in adult *hGFAP-Cre*;*Kdm6b^F/F^*mice

In open field testing, locomotor activity was not impaired in *hGFAP-Cre;Kdm6b^F/F^* mice (**Figure 2—figure supplement 1A, B**). To investigate hippocampal-dependent learning and memory, we used the contextual fear conditioning paradigm as described (Villeda et al., 2014). During fear conditioning training, baseline freezing in *hGFAP-Cre;Kdm6b^F/F^* and littermate controls was not different (**Figure 2A**). However, *hGFAP-Cre;Kdm6b^F/F^* mice had profoundly decreased freezing in contextual memory testing (**Figure 2B**), indicating a defect in hippocampal-dependent behavior. In cued memory testing, hGFAP-Cre;*Kdm6b^F/F^* mice were not different from controls (**Figure 2C, D**), suggesting that non-hippocampal dependent memory was not affected. These results indicate that loss of *Kdm6b* in the developing DG produces defects in hippocampal dependent behaviors.

**Figure 2.**
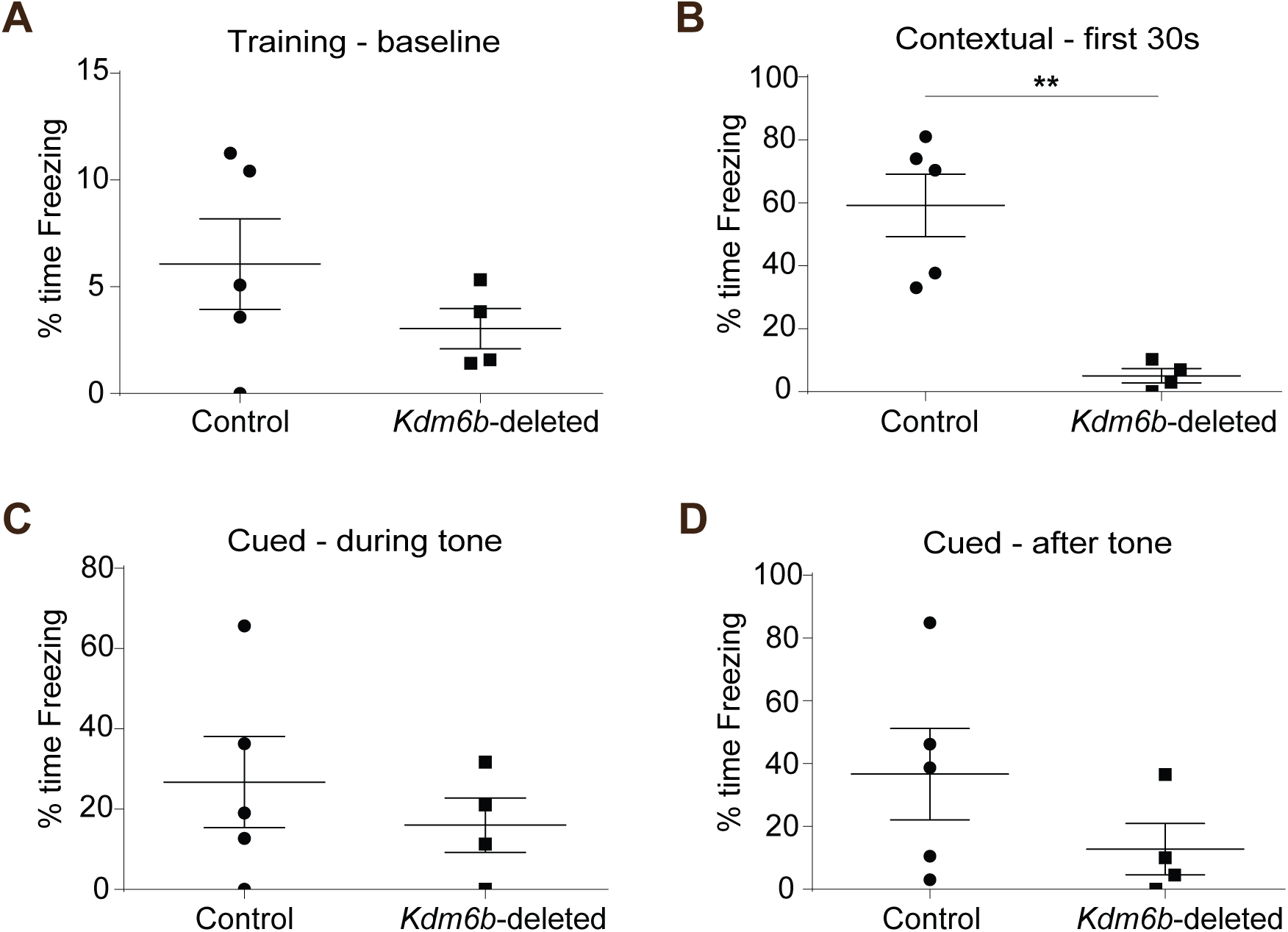
Adult *hGFAP-Cre;Kdm6b^F/F^* mice display defective hippocampal-dependent behavior. **A**, Baseline freezing during fear conditioning training. **B**, Hippocampal-dependent learning and memory assessed in control and *hGFAP-Cre;Kdm6b^F/F^* mice by contextual memory testing. **C-D**, Non-hippocampal dependent learning and memory assessed in control and *hGFAP-Cre;Kdm6b^F/F^* mice by cued memory test. Two-tailed unpaired t test (** = p < 0.01)

### *Kdm6b-*deletion results in a loss of DG NSCs

During the first postnatal week, the DG becomes populated with SOX2+, PAX6+ precursor cells that later give rise to the adult NSC population (Sugiyama et al., 2013; Lugert et al., 2010). In *hGFAP-Cre;Kdm6b^F/F^* mice, there were fewer SOX2+, PAX6+ DG precursor cells at P0 and P7 as compared to controls (**Figure 3A, B**). By P15, DG precursor cells take on a radial morphology characteristic of adult SGZ NSCs (Nicola et al., 2015), and there were fewer radial SOX2+, PAX6+, GFAP+ NSCs in P15 *hGFAP-Cre;Kdm6b^F/F^* mice (**Figure 3—figure supplement 1**). In adulthood, SGZ NSCs express NESTIN and GFAP and extend radial processes into the GCL (Bonaguidi et al., 2011; Kempermann et al., 2015). In adult control mice, we observed NESTIN+, GFAP+ cells in the SGZ with radial processes, corresponding to the SGZ NSC population (**Figure 3C-E**). In stark contrast, in *hGFAP-Cre;Kdm6b^F/F^* mice (n = 10 mutants, 10 controls), we did not identify any cells with these morphological and immunohistochemical characteristics, suggesting the absence of SGZ NSCs in the mutants (**Figure 3F-H**). Young neuroblasts express Doublecortin (DCX), and while many DCX+ cells were observed in the DG of control mice, essentially no DCX+ cells were detected in the DG of *hGFAP-Cre;Kdm6b^F/F^*mice (**Figure 3I, J).** Thus, without *Kdm6b*, the postnatal mouse DG does not establish normal numbers of SGZ NSC, and by adulthood, DG neurogenesis fails.

**Figure 3.**
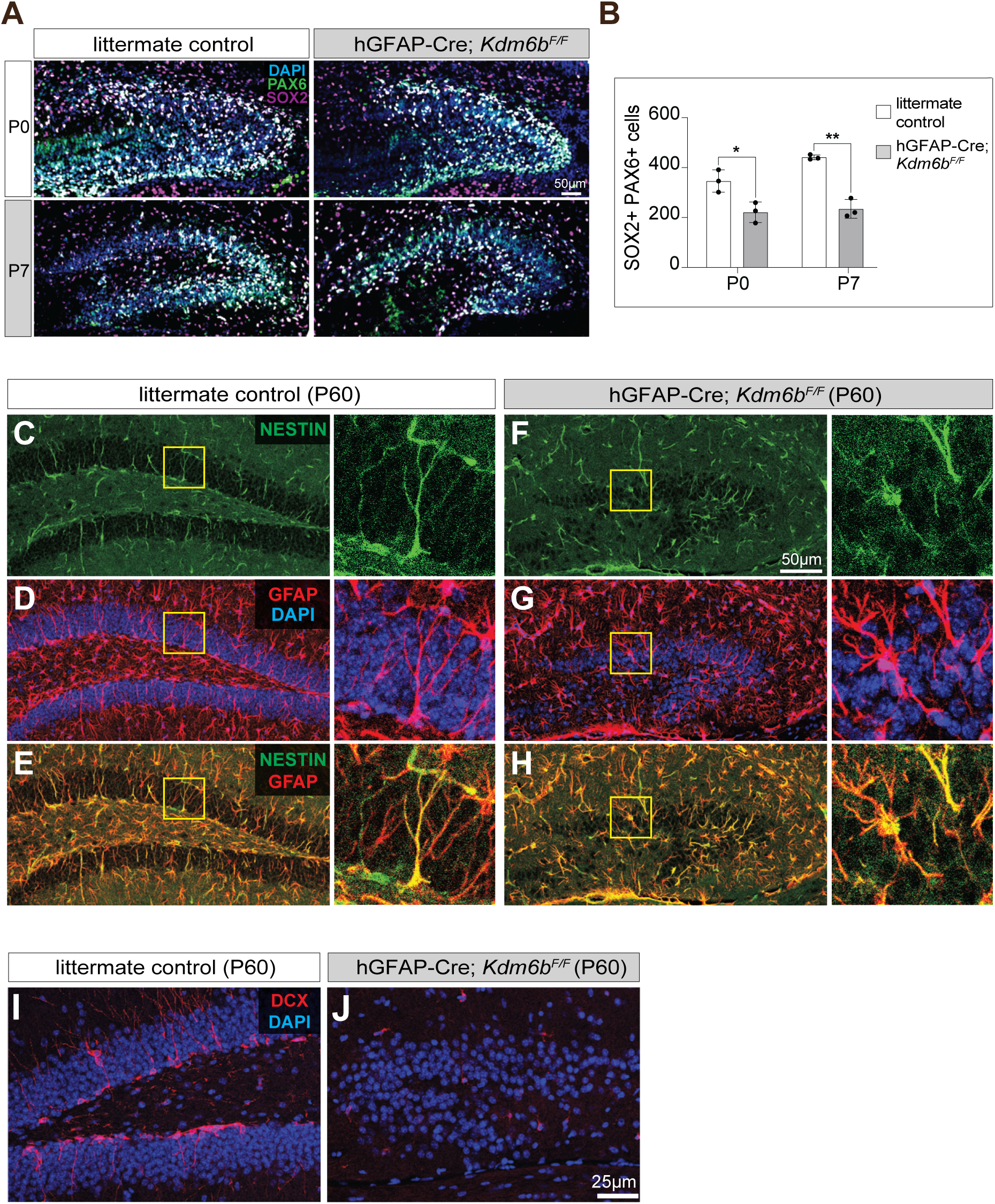
Reduction of NSCs in the developing DG of *hGFAP-Cre;Kdm6b*^F/F^ mice. **A**, IHC for PAX6 (green), SOX2 (purple), and DAPI (blue) in coronal sections of developing DG in control and *hGFAP-Cre;Kdm6b^F/F^*mice at P0 and P7. **B**, Quantification of DG neural precursor/NSCs in control and *hGFAP-Cre;Kdm6b^F/F^* mice at P0 and P7 (n = 3 each), two-tailed unpaired t test (* = p < 0.05, ** = p < 0.01). **C-H**, IHC for NESTIN (green), GFAP (red), and DAPI (blue) in coronal DG sections of P60 control (C-E) and *hGFAP-Cre;Kdm6b^F/F^*mice (F-H). **I-J,** IHC for DCX (red) and DAPI (blue) in coronal DG sections of P60 control (I) and *hGFAP-Cre;Kdm6b^F/F^* mice (J).

Considering the reduced SGZ NSC population in *hGFAP-Cre;Kdm6b^F/F^*mice, we tested for potential changes in cell death by cleaved Caspase3 (Casp3) IHC. However, we did not observe an increase in activated Casp3+ cells in the DG of P0 and P7 *hGFAP-Cre;Kdm6b^F/F^* mice (**Figure 3—figure supplement 2**). Similarly, previous studies have not shown increased cell death with *Kdm6b* deficiency in mouse postnatal subventricular zone NSCs (Park et al., 2014), retinal progenitors (Iida et al., 2014), developing medulla (Burgold et al., 2012) and ESCs (Burgold et al., 2008). The loss of DG NSCs, therefore, does not appear to stem from an increase in cell death.

### *Kdm6b* deletion leads to premature differentiation of embryonic DG precursor cells

Depletion of the SGZ NSC population in the *Kdm6b-*deleted mice at postnatal age could result from premature differentiation of the NSCs during embryonic development. At E15.5, the number of SOX2+ neural progenitor cells (NPCs) as well as the number of SOX2+, EdU+ proliferative cells were not significantly different between *hGFAP-Cre;Kdm6b^F/F^*and control mice (**Figure 4—figure supplement 1**). We injected embryos (*Kdm6b*-deleted vs control) with the cell cycle marker 5-bromo-2-deoxyuridine (BrdU) at E15.5 and subsequently analyzed the DG for BrdU+ cells 2 days and 4 days post injection (dpi) (**Figure 4A**). At E17.5 (2dpi), DG precursor cells start to migrate away from the DNE, forming the dentate migratory stream (DMS) to where the DG blades will develop. (**Figure 4A**). To examine the intermediate progenitor cell (IPC) population, we co-immunostained sections for the IPC marker TBR2. In *Kdm6b*-deleted brains, the number of BrdU+, TBR2+ IPCs was increased compared to littermate control in the stream of migrating precursor cells (**Figure 4B, C**). The increase in number of BrdU+, TBR2+ IPCs was also found in the developing DG blades, accompanied by a decrease in BrdU+, SOX2+ NSCs (**Figure. 4B, D**). At P0.5 (4dpi), this increased number of IPCs differentiated into BrdU+, TBR2+, PROX1+ young neuroblasts, suggesting that excess BrdU+ TBR2+ IPCs are able to further mature into DG neurons in *Kdm6b-*deleted brain (**Figure 5A-C**). These results indicate that SGZ precursor cells prematurely differentiate into neurons in the absence of *Kdm6b*, in accordance with the observed reduction of NSCs in the SGZ of *Kdm6b*-deleted brain (**Figure 5D, E**).

**Figure 4.**
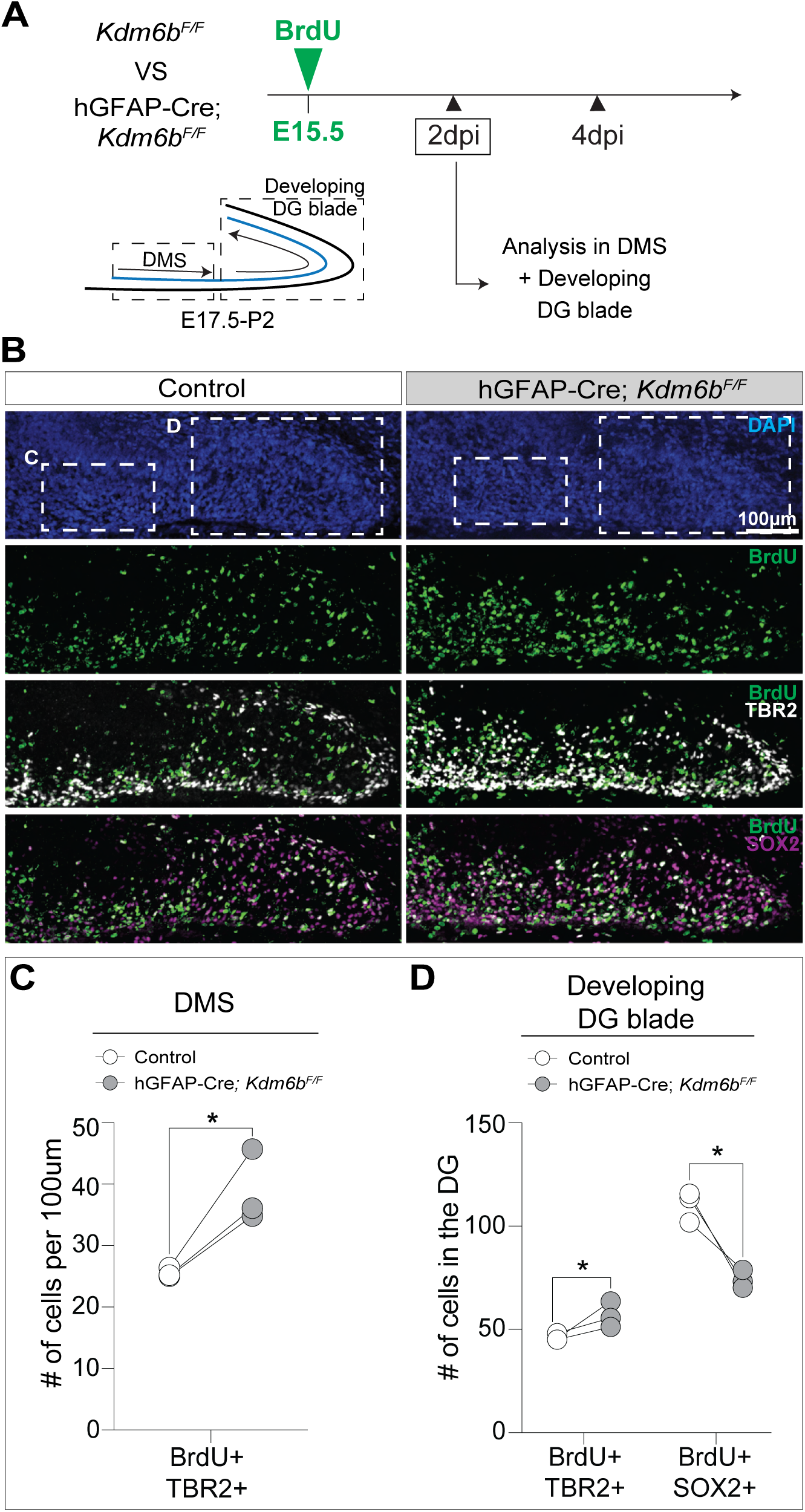
*Kdm6b-*deletion leads to premature differentiation of embryonic DG precursor cells. **A**, Schematic experimental design and coronal section showing the developing dentate gyrus; boxes indicate regions of quantification for (C, D). **B,** IHC for BrdU (green), TBR2 (white), and SOX2 (purple) with DAPI (blue) in coronal sections of DG in control and *hGFAP-Cre;Kdm6b^F/F^* mice at E17.5. **C**, Quantification of BrdU+ TBR2+ cells in dentate migratory stream (DMS) of control and *hGFAP-Cre;Kdm6b^F/F^* mice, two-tailed unpaired t test. **D**, Quantification of BrdU+ TBR2+ and BrdU+ SOX2+ cells in developing DG blades of control and *hGFAP-Cre;Kdm6b^F/F^* mice, two-tailed unpaired t test (* = p < 0.05).

**Figure 5.**
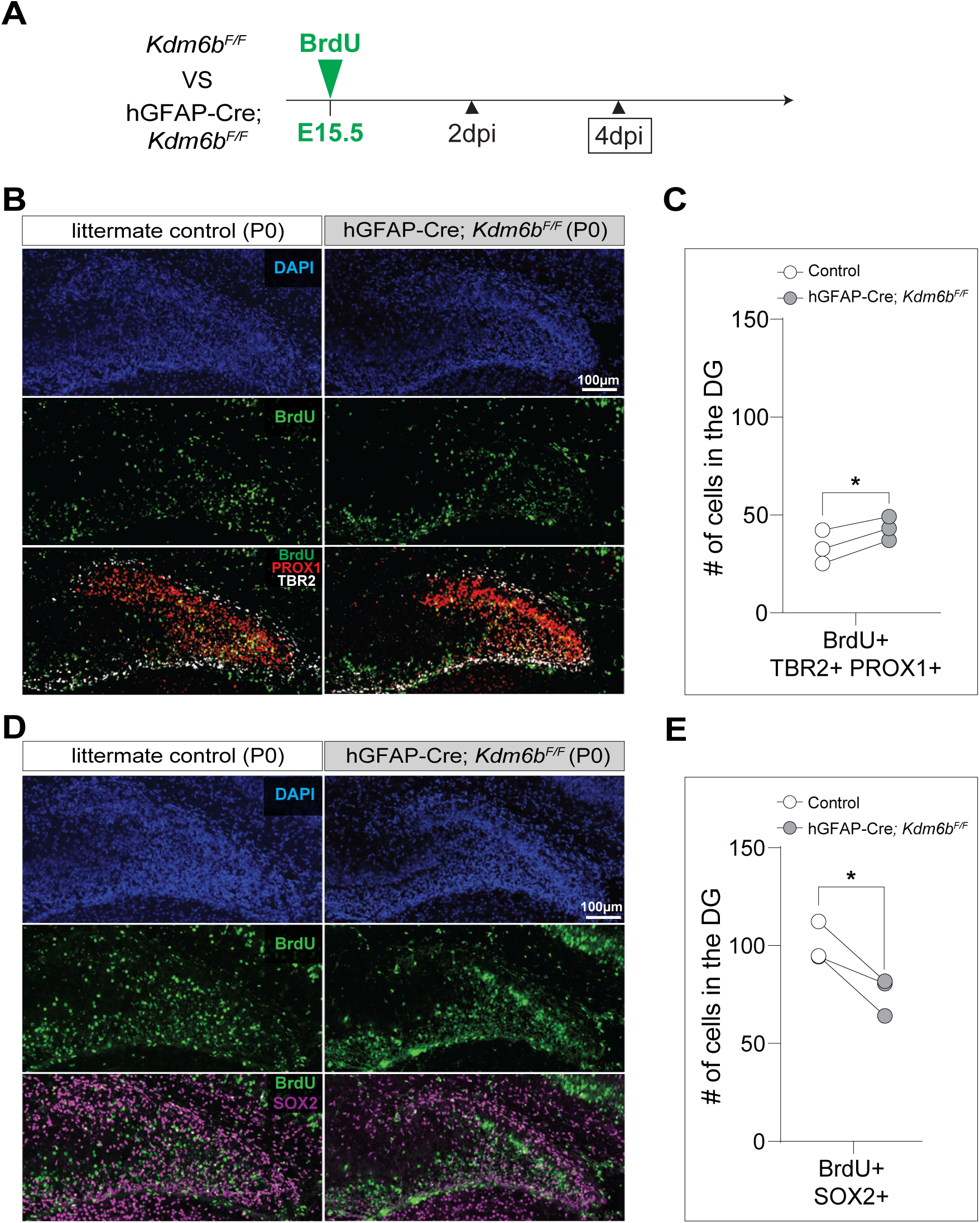
Excess IPCs in *Kd6mb*-deleted mice further mature into DG neurons. **A**, Schematic experimental design. **B**, IHC for BrdU (green), PROX1 (red), TBR2 (white), and DAPI (blue) in coronal sections of DG in control and *hGFAP-Cre;Kdm6b^F/F^*mice at P0.5. **C**, Quantification of (B), two-tailed paired t test (* = p < 0.05). **D**. IHC for BrdU (green), SOX2 (purple), and DAPI (blue) in coronal sections of DG in control and *hGFAP-Cre;Kdm6b^F/F^* mice at P0.5. **E**, Quantification of (D), two-tailed paired t test (* = p < 0.05).

### *Kdm6b*-deleted DG NSCs have an impaired NSC maintenance gene signature

To investigate the specific cell types within the developing DG and aspects of their cell state (*e.g,* stemness, quiescence and proliferation), we performed droplet-based single-cell RNA sequencing (scRNA-seq) on micro-dissected DG from P2 control and *hGFAP-Cre;Kdm6b^F/F^*mice (n=2 per group) (**Figure 6A**). Unsupervised clustering of cells with high-quality transcriptomes (3163 and 2077 DG cells from control and hGFAP-Cre*;Kdm6b^F/F^*animals, respectively) revealed 16 transcriptionally distinct cell clusters, one of which corresponded to NSCs (**Figure 6B**; **Figure 6—figure supplement 1A-D**). *Kdm6b*-deletion was confirmed by examining read alignment data. The NSC cluster (∼13% of the population for both control and *Kdm6b*-deleted DG) co-expressed *Hopx*, *Sox2*, *Gfap*, and *Pax6* (**Figure 6C)**.

**Figure 6.**
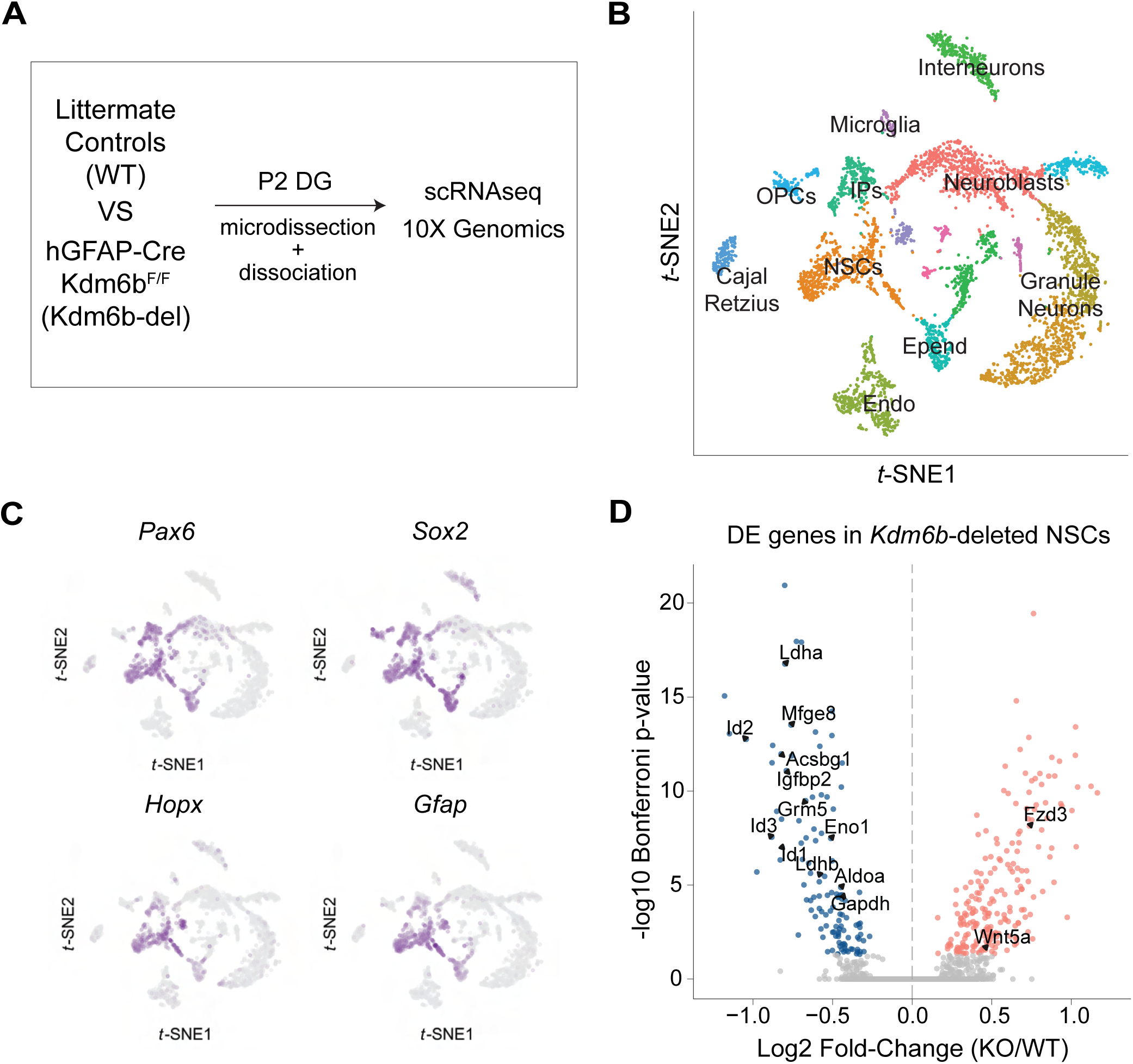
Single cell RNA-seq reveals disrupted stem cell maintenance gene signature in *Kdm6b*-deleted NSCs. **A**, Schematic of single cell RNA-seq experiment design. **B,** t-SNE plot of DG cells from *hGFAP-Cre;Kdm6b^F/F^* mice and control mice labeled with corresponding cell type identity. **C,** t-SNE plot of DG cells with specific marker expression. **D,** Volcano plot of differentially expressed genes in NSCs.

Through differential gene expression analysis of NSCs from hGFAP-Cre*;Kdm6b^F/F^* mice and littermate controls, we identified 337 differentially expressed (DE) genes (119 and 218, down-regulated and up-regulated in *Kdm6b*-deleted NSCs, respectively) at a Bonferroni adjusted *p*-value < 0.05 (**Figure 6D**). In a previous single-cell RNA-seq study of adult DG NSCs, the activation of NSCs for neurogenesis is accompanied by a decline in gene expression for glycolysis, fatty acid degradation, and drug metabolism pathways (Shin et al., 2015). In *Kdm6b*-deleted NSCs, down-regulated genes were enriched for multiple Gene Ontology (GO) terms related to glycolysis (**Figure 6—figure supplement 1E**), similar to the progressive gene expression changes during normal NSC activation (Shin et al., 2015). *Acsbg1*, a gene involved in the first step of fatty acid β-oxidation, is down-regulated during NSC activation (Shin et al., 2015) and was also decreased in *Kdm6b*-deleted NSCs. Drug metabolism is another biological process down-regulated in activated NSCs (Shin et al., 2015), and *Hspd1*, *Pgk1*, *Pgam1*, *Eno1*, *Gpi1* – genes with cell-intrinsic roles in drug metabolism – were all down-regulated in *Kdm6b*-deleted NSCs. On the other hand, some genes upregulated in *Kdm6b*-deleted NSCs (*Wnt5a, Fzd3*) have been reported to promote neuronal differentiation (Park et al., 2018; Arredondo et al., 2020). These data suggest that *Kdm6b*-deletion causes metabolic changes that correspond to a shift away from NSC self-renewal and progression towards neurogenesis.

Other genes implicated in NSC self-renewal including insulin growth factor binding protein 2 (*Igfbp2*), milk fat globule-EGF factor 8 (*Mfge8*), and inhibitors of DNA binding and cell differentiation (*Id1, Id2, Id3*), were down-regulated in *Kdm6b*-deleted NSCs (**Figure 6D**). *Igfbp2* promotes self-renewal and proliferation of NSCs, with knockdown of the gene resulting in precocious neuronal differentiation (Shen et al., 2018). *Mfge8* is also essential for NSC maintenance, as deletion of *Mfge8* leads to overactivation and subsequent depletion of DG NSCs (Zhou et al., 2018). Similarly, *Id1*, *Id2*, and *Id3* have been previously shown to be essential for embryonic brain NSC maintenance, with knockout or knockdown causing precocious neuronal differentiation (Niola et al., 2012). Together, these data indicate that *Kdm6b*-deletion results in a transcriptomic signature that underlies their defective NSC maintenance.

### *Kdm6b* deletion increases H3K27me3 levels at regulatory elements

H3K27me3 is a chromatin modification that correlates with transcriptional repression, and KDM6B is a H3K27me3 demethylase (Agger et al., 2007; Burchfield et al., 2015; Hong et al., 2007). We performed Cleavage Under Targets & Release Using Nuclease (CUT&RUN) analysis on P2 DG to assess H3K27me3 levels in *Kdm6b*-deleted vs. littermate control cells (**Figure 7A)**. In WT cells, low levels of K27me3 levels at the transcriptional start site (TSS) correlate with higher levels of gene expression (**Figure 7B**). Genes down-regulated in *Kdm6b*-deleted DG were assessed by pooling scRNA-seq data from P2 DG, and of the 119 genes down-regulated in the NSC population, 94 (79.0%) were also at decreased levels across all DG cell types including 14 (out of 16, 87.5%) genes with roles in NSC maintenance (**Figure 7C**). 11 out of 16 (68.8%) of the downregulated NSC maintenance genes (*e.g., Id* genes) exhibited increased levels of H3K27me3 at the TSS in *Kdm6b*-deleted DG cells (**Figure 7D-F**). These data indicate that the histone demethylase activity of KDM6B corresponds to the chromatin state that underlies NSC maintenance.

**Figure 7.**
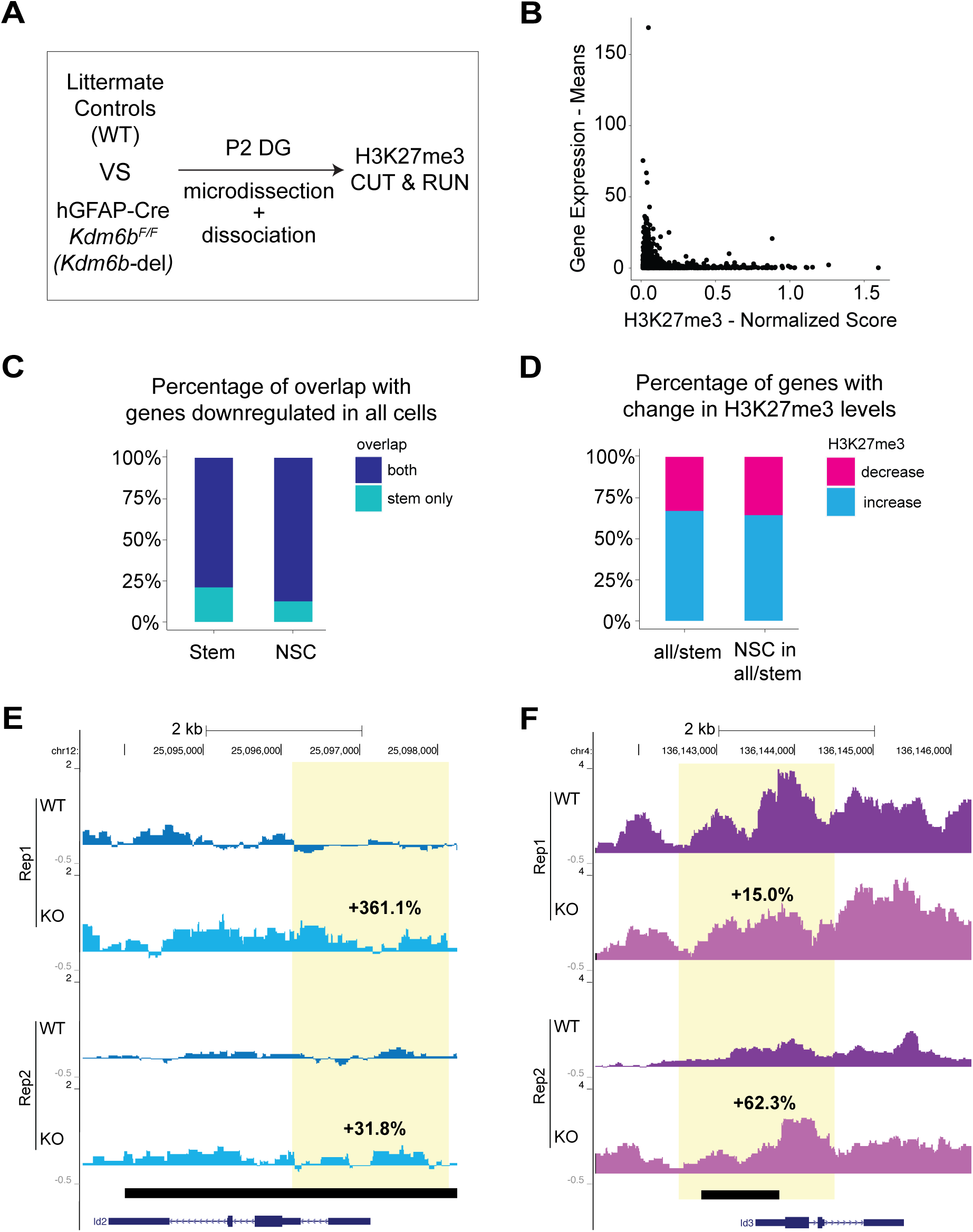
*Kdm6b*-deletion increases H3K27me3 levels at regulatory elements. **A**, Schematic of CUT&RUN experiment design. **B,** Correlation plot of H3K27me3 enrichment and gene expression in control mice. **C**, Bar on left depicts the overlap between genes downregulated in stem cells and in all cells pooled together. Bar on right depicts the overlap between NSC maintenance genes and genes downregulated in all cells pooled. **D**, Bars depict changes in H3K27me3 enrichment for overlapping genes shown by the left and right bars in (C), respectively. **E-F**, H3K27me3 signal over the *Id2* locus (E) and *Id3* locus (F) in cells from control and *Kdm6b*-deleted DG. Yellow column indicates +/1kb of the transcriptional start site (TSS). Black bar indicates regions with statistically significant differences in H3K27me3 enrichment. Two biological replicates are shown.

### *Kdm6b* is required for the maintenance of adult hippocampal NSCs

After becoming established in the DG during postnatal development, the NSC population is maintained throughout adult life. To investigate the role of *Kdm6b* in NSC maintenance, we acutely deleted *Kdm6b* in adult mouse DG NSCs with the tamoxifen (TAM)-inducible Nestin-Cre^ERT2^ transgene (Lagace et al., 2007). To follow the fate of cells that had undergone recombination, we used the Ai14 Cre-reporter transgene, which expresses tdTomato after excision of a “floxed-stop” cassette (Madisen et al., 2010). Nestin-Cre^ERT2^; *Kdm6b^F/F^*;Ai14 and Nestin-Cre^ERT2^ ;*Kdm6b^F/+^*;Ai14 (littermate controls) mice were administered tamoxifen (TAM) for 5 days starting at P60 (**Figure 8A**). At 1 day post injection (dpi) of TAM administration, the number of tdTomato+ cells in the DG of Nestin-Cre^ERT2^; *Kdm6b^F/F^* mice was not different as compared to littermate controls (**Figure 8B**). However, at 10dpi, we observed a 28% increase in tdTomato+, Tbr2+ intermediate progenitors and a 43% increase in tdTomato+, DCX+, Tbr2+ neuroblasts in Nestin-Cre^ERT2^;*Kdm6b^F/F^*;Ai14 mice (**Figure 8C, D**). To investigate whether this precocious neuronal differentiation is accompanied by a loss of NSCs, we quantified the DG NSC population of animals at 30dpi. In Nestin-Cre^ERT2^; *Kdm6b^F/F^*;Ai14 mice, there were 25% fewer tdTomato+ adult NSCs – defined by the expression of SOX2 and radial cell morphology – as compared to littermate controls (**Figure 8E, F**). Thus, in adult mice, *Kdm6b*-deletion causes precocious neuronal differentiation and depletion of the NSC population.

**Figure 8.**
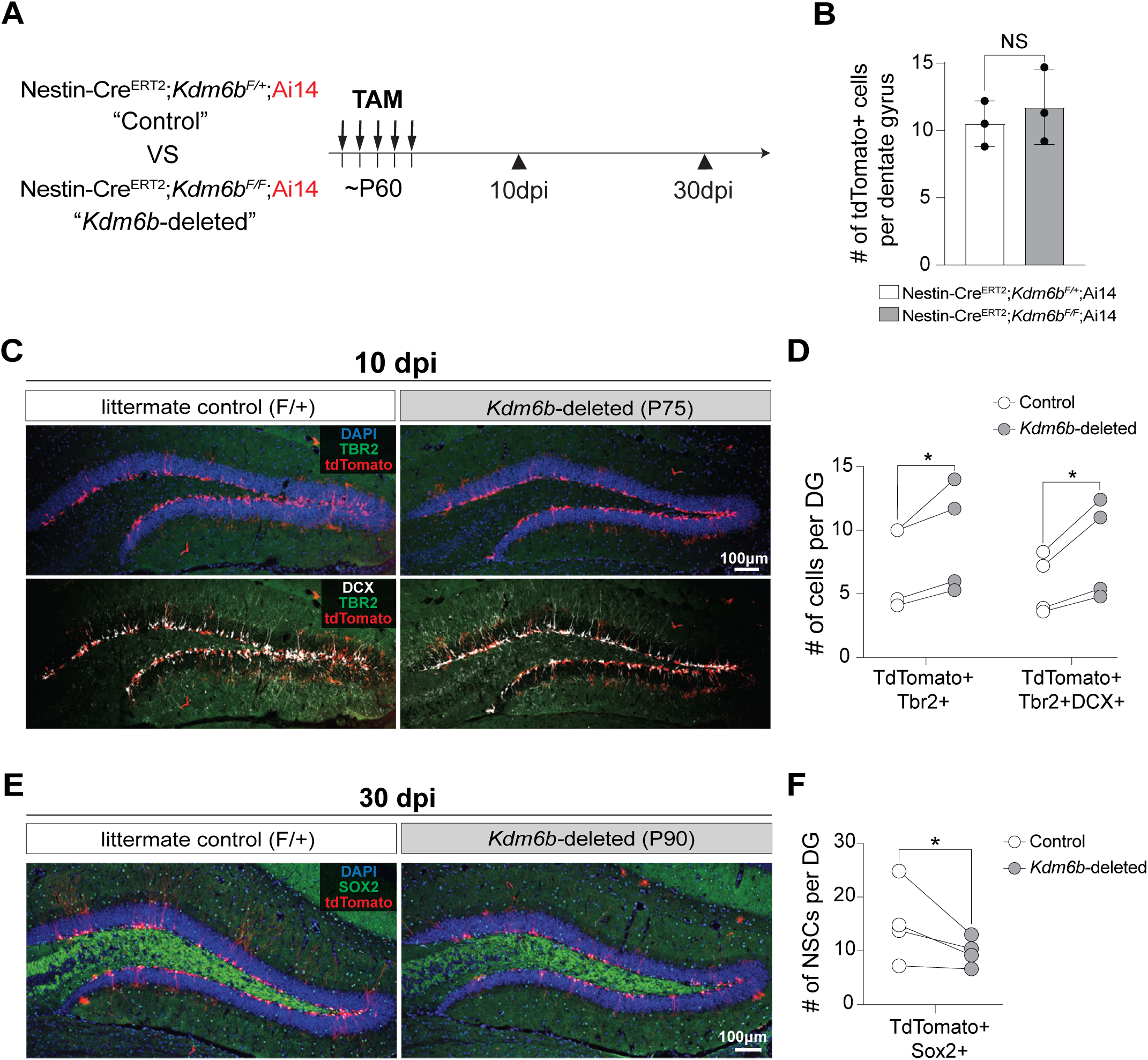
*Kdm6b* is required for maintenance of adult SGZ NSCs. **A**, Schematic of experimental design. **B,** Quantification of number of tdTomato+ cells at the end of TAM administration comparing Nestin-Cre^ERT2^;*Kdm6b^F/F^*;Ai14 to control. Two-tailed unpaired t test (NS = not significant). **C**, IHC for tdTomato (red), TBR2 (green), DCX (white), and DAPI (blue) in coronal sections of DG in control and Nestin-Cre^ERT2^; *Kdm6b^F/F^*;Ai14 mice at 10dpi. **D**, Quantification of (B), two-tailed paired t test (* = p < 0.05). **E**, IHC for tdTomato (red), SOX2 (green), and DAPI (blue) in coronal sections of DG in control and Nestin-Cre^ERT2^; *Kdm6b^F/F^*;Ai14 mice at 30dpi. **F**, Quantification of (D), two-tailed paired t test (* = p < 0.05).

## DISCUSSION

Towards the end of brain development, most neural precursors exit the cell cycle and terminally differentiate into postmitotic neurons and glia, effectively leading to a depletion of stem cells. However, a small population of stem cells takes up residence in the hippocampal DG of the adult mouse and continues to produce neurons for the duration of the animal’s life. The molecular mechanisms that underlie how NSCs become established and maintained in the adult brain are still not well understood. In this report, we demonstrate that *Kdm6b* is essential for both the establishment and long-term maintenance of NSCs in the mouse hippocampal DG. Below, we discuss our results in the context of adult neurogenesis, transcriptional mechanisms underlying NSC maintenance, and potential relevance to human cognitive ability.

In the mouse, adult NSC populations become established in two different brain regions: the ventricular-subventricular zone (V-SVZ) and the DG. Although both populations of adult NSCs are astrocyte-like cells that arise from embryonic brain radial glia, the deletion of *Kdm6b* impacts the two groups differently. In the V-SVZ, NSCs populate the walls of the lateral ventricle regardless of the presence of *Kdm6b*, indicating that this chromatin regulator is not required for the establishment of the population (Park et al., 2014). Without *Kdm6b*, the number of V-SVZ NSCs as defined by their gene expression and unique morphology increased 3- to 4-fold as compared to littermate controls (Park et al., 2014). In contrast, without *Kdm6b*, the DG NSC population never properly became established. Instead, the proliferating *Kdm6b*-deleted DG precursor cells failed to acquire the typical morphology and location of DG NSCs, ultimately producing a DG essentially devoid of adult neurogenesis.

The small and hypocellular DG of adult *hGFAP*-Cre; *Kdm6b^fl/fl^*mice was a result of precocious neuronal differentiation leading to depletion of postnatal DG neural precursors. In the V-SVZ, NSC lacking *Kdm6b* instead become “stalled” during neuronal differentiation, failing to fully activate the expression of key neurogenic transcription factors such as *Dlx2* (Park et al., 2014). Thus, while *Kdm6b* is critical for NSC establishment and restraint of neurogenesis in the DG, *Kdm6b* appears to support the neuronal specification of NSCs in the V-SVZ, highlighting the distinct roles that KDM6B plays in regulating the two NSC populations.

Consistent with the observed precocious DG NSC differentiation in mutant mice, there was a decrease in expression of genes known to promote NSC maintenance in our transcriptomic analysis of *Kdm6b*-deleted DG NSCs. This group of genes included members of the Id family, which encodes proteins that inhibit the function of basic HLH transcription factors involved in cell differentiation (Perk et al., 2005). Triple knockout of *Id1-3* increases neuronal differentiation in mice NSCs, while there is a loss of NSCs *in vivo* in the postnatal V-SVZ (Niola et al., 2012). On a more global level, our transcriptomic analysis of *Kdm6b*-deleted DG NSCs indicated a shift away from glycolytic metabolism, similar to that seen in the activation of quiescent DG NSCs (Shin et al., 2015). These data support a model wherein the inability to maintain postnatal DG neural precursors results in the failure to establish the adult NSC population in the mouse DG.

Acute deletion of *Kdm6b* also led to precocious differentiation of adult DG NSCs, suggesting that *Kdm6b* continues to maintain the NSC population throughout life. Without successful establishment and maintenance of an adult DG NSC population, hippocampal neurogenesis cannot persist. For instance, deficiency in Notch signaling (from the *Notch2* membrane receptor to the *Hes1* transcription factor) in mouse NSC populations causes precocious neuronal differentiation and depletion of the NSC pool, resulting in a loss of adult neurogenesis (Engler et al., 2018; Lampada and Taylor, 2023). Several chromatin regulators, which play important roles in orchestrating broad transcriptional programs, are known to be necessary for adult mouse DG neurogenesis (Hsieh and Eisch, 2010). Still, their link to NSC maintenance remains unclear. Examples include CHD7, required for DG neuronal differentiation but dispensable for NSC maintenance (Feng et al., 2013), and BRG1, with its deletion resulting in an initial decrease of adult DG neuron production but having no visible impact on the NSC population (Petrik et al., 2015). Although EZH2, a Polycomb factor that in principle opposes KDM6B by catalyzing H3K27me3, is required for the proliferation of neural progenitor cells, it does not appear to play a role in the quiescent NSC population (Zhang et al., 2014). A truncating mutation of *Kmt2d* in mice was notable for leading to a reduced Nestin positive NSC population, but a reduction in IPCs and neuroblasts was also observed (Carosso et al., 2019). In contrast to these prior studies of other chromatin regulators, the phenotype of *Kdm6b*-deletion is more consistent with a precocious activation of neurogenesis from the NSC pool, resulting in its depletion.

In our CUT&RUN analysis, the majority – but not all – of downregulated NSC maintenance genes exhibited increased levels of H3K27me3 at their promoter regions with *Kdm6b*-deletion, suggesting that KDM6B-mediated regulation of the NSC transcriptome is dependent on both its catalytic demethylase activity and demethylase-independent activity. KDM6B’s mechanism of action is often presumed to be gene activation via removal of H3K27me3, a mark correlated with transcriptional repression. Correspondingly, prior studies have shown that *Kdm6b*-deficiency leads to enrichment of H3K27me3 at downregulated genes (Agger et al., 2009; Park et al., 2014; Ramesh et al., 2023). However, KDM6B also has demethylase-independent functions. For example, KDM6B can mediate chromatin remodeling via its interactions with the T-box transcription factor T-bet and the SWI/SNF complex (Miller et al., 2010) and also coordinate the formation of phase-separated condensates to drive enhancer-cluster assembly and gene activation (Vicioso-Mantis et al., 2022). Thus, it is possible that the demethylase independent functions of KDM6B contribute to the regulation of DG neurogenesis.

Mutations in *KDM6B* have been consistently linked to deficits in cognitive function (Gao et al., 2022; Wang et al., 2022; Rots et al., 2023), and multiple studies have previously identified *KDM6B* as a risk gene for autism spectrum disorder (ASD) (De Rubeis et al., 2014; Iossifov et al., 2014; Sanders et al., 2015; Satterstrom et al., 2020). In 85 patients with pathogenic *KDM6B* variants, over 90% of individuals were diagnosed with ASD and/or intellectual disability (Rots et al., 2023). In mice, *Kdm6b* haploinsufficiency results in behavioral deficits linked to ASD (Gao et al., 2022). Postnatal deletion of *Kdm6b* in cortical and hippocampal excitatory neurons of mice also impairs memory and learning (Wang et al., 2022). Our *Kdm6b* mutant mice similarly displayed decreased freezing in a fear conditioning test, a behavior associated with impaired hippocampal-dependent memory. Further testing of KDM6B in systems such as human cell lines or organoids could lead to greater understanding of their mechanisms by which KDM6B regulates hippocampal development, and subsequently cognitive ability, in humans.

## MATERIALS AND METHODS

### Mice

Wildtype (C57bl/6J), hGFAP-Cre: Tg(GFAP-Cre)25Mes/J (Zhuo et al., 2001), Nestin-Cre^ERT2^: Tg(Nes-cre/ERT2)KEisc/J (Lagace et al., 2007), and Ai14: Gt(ROSA)26Sor^tm14(CAG-tdTomato)Hze^ (Madisen et al., 2010) mice were obtained from Jackson Laboratory. *Kdm6b^F/F^* mice which contain *Kdm6b* alleles with loxP sites flanking the JmjC catalytic domain were maintained and genotyped as described (Iwamori et al., 2013; Park et al., 2014). To label proliferating cells, BrdU (10 mg/ml solution, 50 μg/gram body weight, Sigma) was injected intraperitoneally. Experiments were performed in accordance to protocols approved by Institutional Animal Care and Use Committee at UCSF. For the purpose of adult fate-tracing, >P60 mice received 5mg of TAM (Sigma) dissolved in 100% corn oil (Sigma) by oral gavage per 30 grams of the body weight once a day for 5 consecutive days.

### BrdU administration

Mice were injected with 5-bromo-2’-deoxyuridine (BrdU, Millipore Sigma) reconstituted in sterile PBS (10mg/ml) intraperitoneally at a dose of 50mg of BrdU per kg in mouse weight.

### Tissue Preparation and Immunohistochemistry

For embryonic to early postnatal timepoints (P0, P7), mouse brains were quickly immersed into 4% paraformaldehyde (PFA) after extraction and fixed at 4°C overnight. The brains were then washed with phosphate-buffered saline (PBS) and stored at 4°C overnight. Afterwards, the brains were incubated in 30% sucrose for 24-36 hours before they were frozen inside the cryo-mold using a 1:1 mixture of 30% sucrose and OCT. Brain sections were collected coronally on the cryostat at 12μm thickness. For P15 to adult timepoints, the mice first underwent intracardiac perfusion (Park et al., 2014) with PBS followed by 4% PFA. The brains were then post-fixed in 4% PFA at 4°C overnight. The following steps prior to sectioning were the same as listed above. Brain sections for these ages were collected coronally on the cryostat at 14μm thickness.

Sections were blocked with with 10% normal goat/donkey serum, 0.3% Triton-X 100, 1% bovine serum albumin, and 0.3M glycine in PBS for 1hr at room temperature. Primary antibodies were diluted in blocking buffer and incubated at 4 °C overnight. For JMJD3 staining, rabbit anti-JMJD3 antibodies (Abgent) were affinity purified using EpiMAX Affinity Purification kit (Abcam) and epitope retrieval was performed with 2N HCl. Fluorescence signal was amplified using TSA Plus fluorescence kit (PerkinElmer). The following primary antibodies were used in this study: rabbit anti-JMJD3 (epitope purified, 1:5, Abgent), rat anti-GFAP (1:500, Invitrogen), guinea pig anti-Doublecortin (1:1000, Millipore), rat anti-BrdU (1:500, Abcam), mouse anti-NeuN (1:500, Chemicon), mouse anti-NESTIN (1:500, Millipore), rabbit anti-Cleaved Caspase 3 (1:250, Covance), goat anti-SOX2 (1:300, Santa Cruz) rabbit anti-PROX1 (1:300, Covance). After washing in PBS, sections were incubated with DAPI (1:1000, Sigma) and Alexa-Fluor secondary antibodies (Invitrogen) diluted in blocking buffer for 2 hours at RT. Samples were mounted with Aqua-poly-mount (Polysciences Inc).

### Microscopic Analysis

Confocal images were obtained using a Leica TCS SP5X and the Leica Stellaris 5. Sections from postnatal mice were imaged at either 10X or 20X magnification with 4096 x 4096 resolution for the Leica TCS SP5X and 2048 x 2048 resolution for the Leica Stellaris 5. Sections from embryonic mice were imaged at 10X with 4096 x 4096 resolution. For *in vivo* DG cell quantification, we used at least 3 animals per group. From each animal, at least three adjacent brain sections were imaged within the slide and used to quantify resulting in a total of 6 (or more) hippocampi. For embryonic to early postnatal age, each brain sections were 84μm apart in the anterior-posterior axis. For later postnatal to adult, sections were 98μm apart in the AP axis. Image processing and quantifications were completed in ImageJ. Statistical tests of significance were performed using t-Test in GraphPad Prism 5.

### *In Situ* Hybridization (ISH)

ISH on brain tissue was performed as previously described (Wallace and Raff, 1999) with DIG-labeled RNA probe designed against JmjC domain of *Kdm6b*. ISH images were obtained using a DMI4000B microscope (Leica).

### Open Field

Paradigm follows previously published protocol (Wheatley et al., 2019). Mice were placed in the center of an open 40-cm × 40-cm square chamber (Kinder Scientific) with no cues or stimuli and allowed to move freely for 10 min. Infrared photobeam breaks were recorded and movement metrics analyzed by MotorMonitor software (Kinder Scientific).

### Contextual fear conditioning

Paradigm follows previously published protocol (Villeda et al., 2014).

### Single-cell RNA-seq Analysis

DG were microdissected from P2 *hGFAP-Cre;Kdm6b^F/F^* mice and littermate controls (2 animals per condition) and dissociated into single cells using the Worthington Papain Dissociation system following all manufacturer protocol steps including ovomucoid gradient. Single-cell libraries were generated using the 10x Genomics Chromium Single Cell 3’ Assay with a targeted cell recovery of 5000 (2500 cells per condition). Libraries were sequenced to a mean depth of approximately 55,000 reads/cell and processed through cellranger 3.0.2 (10x Genomics) with a raw recovery of 5847 cells (3555 WT and 2292 KO). All subsequent processing and analyses were performed using the Seurat v3 R package (Butler et al., 2018). Quality filtering was performed on the basis mitochondrial and ribosomal content before log normalization and scaling. WT and KO datasets were then integrated using canonical correlation analysis and the integrated dataset was used for dimensionality reduction by principal components analysis (PCA) and *t*-distributed stochastic neighbor embedding (*t*-SNE). Cluster analysis on the integrated was conducted using the Louvain algorithm on the shared nearest neighbor network, with resolution set to 0.4. Major cell types were assigned to clusters using canonical markers from the experimental literature as well as other published scRNA-seq datasets of the rodent hippocampus. Differential expression testing was performed between WT and KO cells within pertinent clusters using the likelihood-ratio test for single-cell gene expression (“bimod” setting) and p-values were adjusted using the Bonferroni correction.

### CUT&RUN

Brains were extracted from P2 *hGFAP-Cre;Kdm6b^F/F^* mice and littermate controls (3 animals per condition) and sectioned on the vibratome at 400nm thickness. The DG was then microdissected out and dissociated into single cells using the Worthington Papain Dissociation system. 15,000-20,000 isolated cells were used for each experiment. CUT&RUN was performed as previously described (Ahanger et al., 2021) with the following antibodies: anti-H3K4me3 (Cell Signaling Technologies, no. 9751, 1:100) and anti-H3K27m3 (Cell Signaling Technologies, no. 9733, 1:100) An IgG control was run in parallel for each experiment (Cell Signaling Technologies, no. 2729, 1:100).

### Library preparation, sequencing and data processing

CUT&RUN libraries were prepared using KAPA HyperPlus with amplification DNA library preparation kit (no. KK8512) as previously described (Ahanger et al., 2021). 150-bp paired-end sequencing of the libraries were then performed on the Illumina HiSeq 4000. Low-quality reads and adaptors were removed with BBduk (BBMap – Bushnell B. – sourceforge.net/projects/bbmap/), and reads were aligned to the mouse (UCSC mm10) genome using bowtie2 (Langmead and Salzberg, 2012) with the following settings: -local-very-sensitive-local-no-unal-no-mixed-nodiscordant -q-phred33 pI 10 -X 700 (Skene and Henikoff, 2017). Bigwig files were made from aligned reads using deepTools (v3.2.0) bamCompare at 10-bp bin size. Differential enrichment analysis of H3K27me3 enrichment between *Kdm6b*-deleted and control DG was performed in biological duplicates using DiffReps (Shen et al., 2013) in “block” mode and with a p-value cutoff of 0.05. These significantly different areas were then intersected with peaks called by MACS2 (Zhang et al., 2008) with the -broad parameter. H3K27me3 enrichment at +/1 kb of the TSS for each gene was calculated via multiBigwigSummary from deepTools.

## Supplemental Figures

**Figure 1—figure supplement 1.**
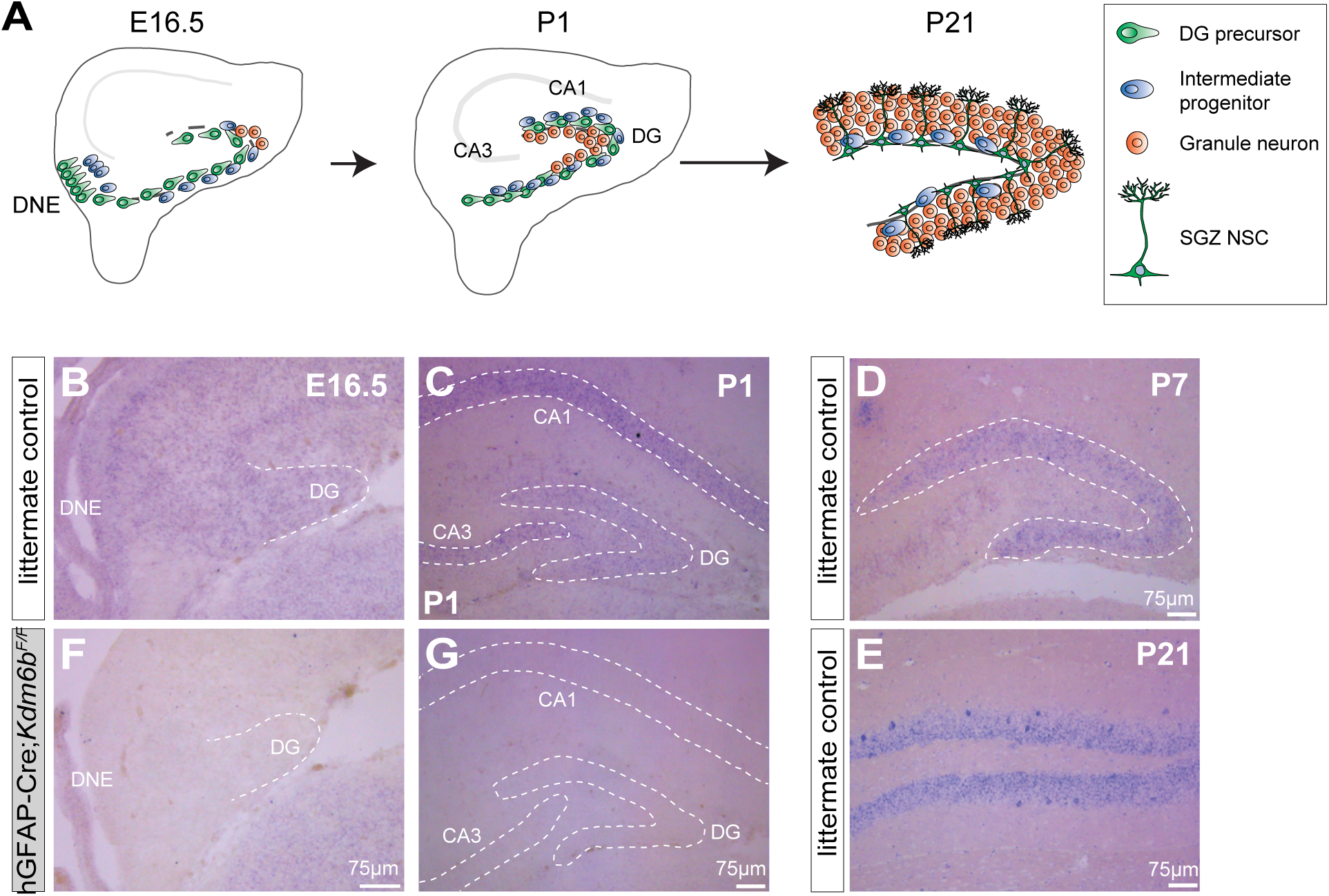
Expression of *Kdm6b* during DG development. **A**, Schematic illustration of DG in development. **B**, ISH for *Kdm6b* in coronal DG sections of control mice at E16.5. **C,** ISH for *Kdm6b* in coronal DG sections of control mice at P1. **D**, ISH for *Kdm6b* in coronal DG sections of control mice at P7. **E,** ISH for *Kdm6b* in coronal DG sections of control mice at P21. **F**, ISH for *Kdm6b* in coronal DG sections of *hGFAP-Cre;Kdm6b^F/F^*mice at E16.5. **G**, ISH for *Kdm6b* in coronal DG sections of *hGFAP-Cre;Kdm6b^F/F^*mice at P1.

**Figure 2—figure supplement 1.**
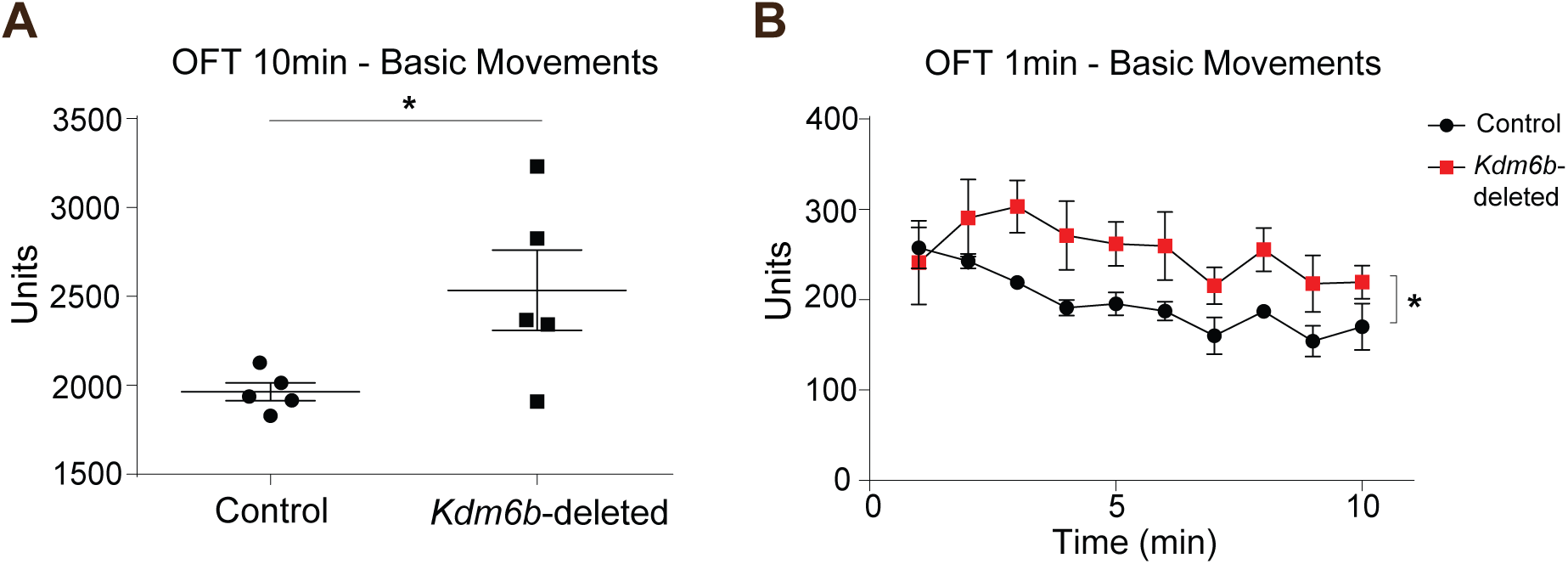
*Kdm6b*-deleted mice do not display locomotive defects. **A**, Quantification of total movement over 10 minutes (measured in units) in the open field paradigm. Two-tailed unpaired t test (* = p < 0.05). **B**, Quantification of locomotion by minute (measured in units) in the open field paradigm. Repeated measures ANOVA with Bonferroni *post-hoc* test (* = p < 0.05).

**Figure 3—figure supplement 1.**
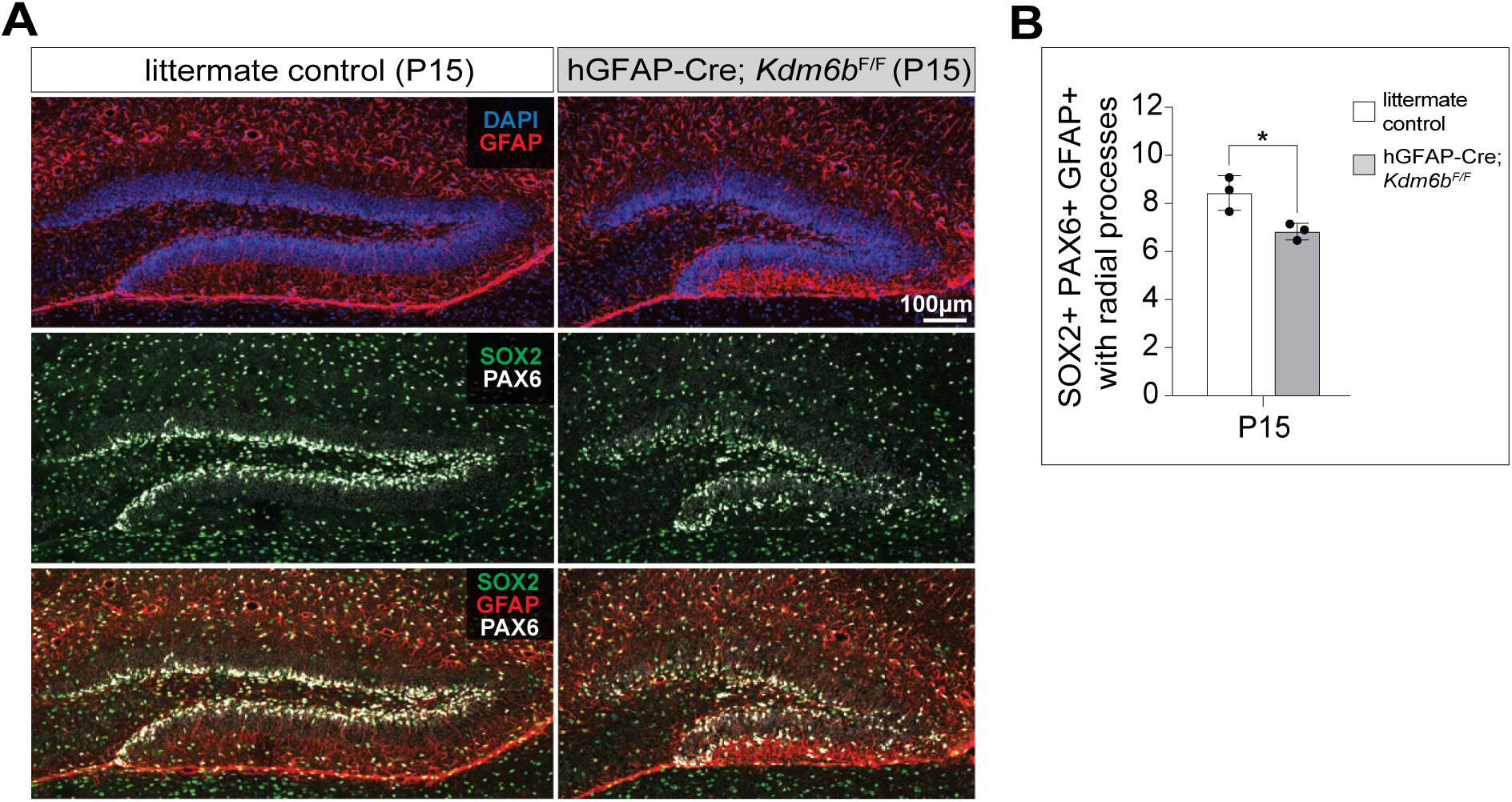
Reduction of NSCs in the DG of *hGFAP-Cre;Kdm6b*^F/F^ mice at P15. **A**, IHC for PAX6 (white), SOX2 (green), GFAP (red), and DAPI (blue) in coronal sections of DG in control and *hGFAP-Cre;Kdm6b^F/F^* mice at P15. **B**, Quantification of DG neural precursor/NSCs in control and *hGFAP-Cre;Kdm6b^F/F^*mice at P15 (n = 3 each), two-tailed unpaired t test (* = p < 0.05).

**Figure 3—figure supplement 2.**
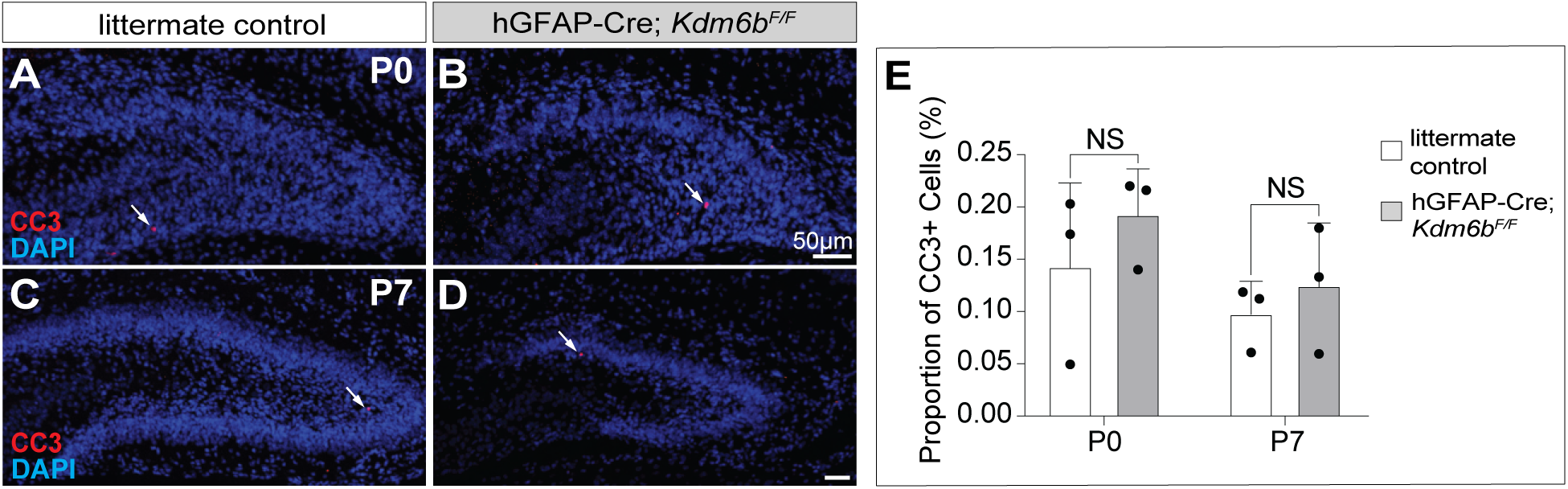
Increased cell death is not observed in *hGFAP-Cre;Kdm6b*^F/F^ mice. **A-B**, IHC for cleaved caspase 3 (CC3) (red) and DAPI (blue) in coronal DG sections of P0 control (A) and *hGFAP-Cre;Kdm6b^F/F^* mice (B). **C-D**, IHC for CC3 (red) and DAPI (blue) in coronal DG sections of P7 control (C) and *hGFAP-Cre;Kdm6b^F/F^* mice (D). **E**, Quantification of (A-D) two-tailed unpaired t test (NS = not significant).

**Figure 4—figure supplement 1.**
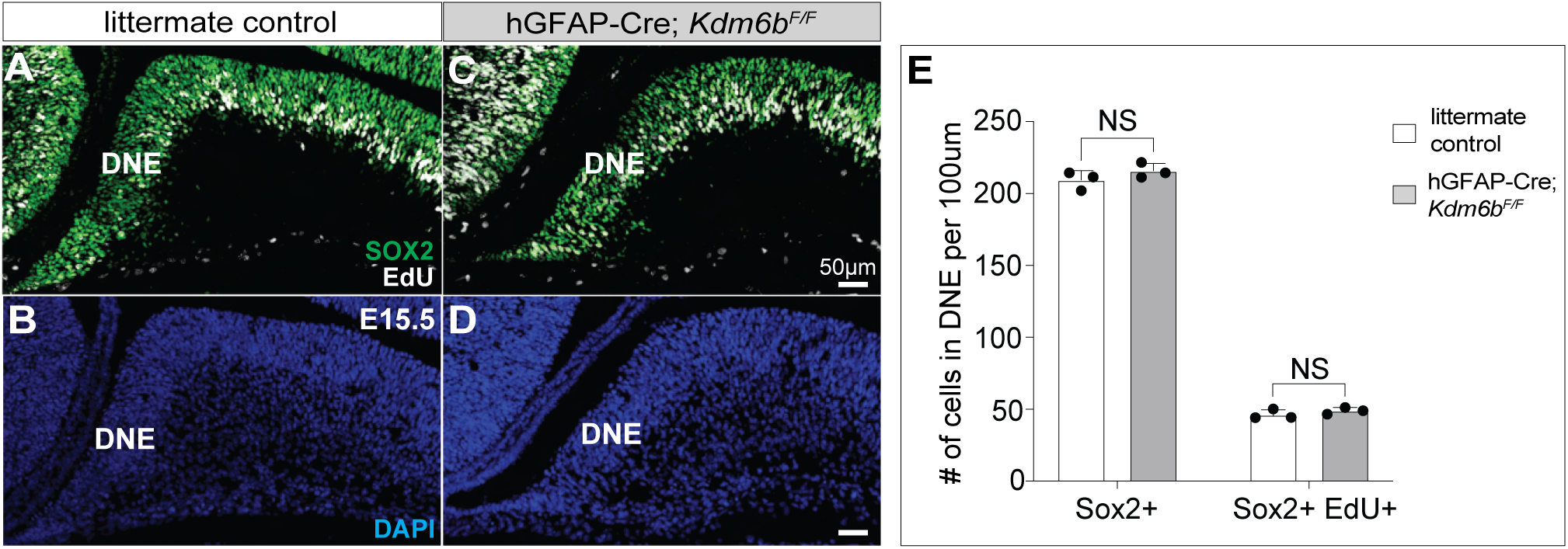
Similar number of SGZ NSC precursor cells at E15.5 between *Kdm6b*-deleted and control brain. **A-D,** IHC for SOX2 (green), EdU (white) and DAPI (blue) in coronal DG sections of E15.5 control (A, B) and *hGFAP-Cre;Kdm6b^F/F^* mice (C, D). **C,** Quantification of (A, C) two-tailed unpaired t test (NS = not significant).

**Figure 6—figure supplement 1.**
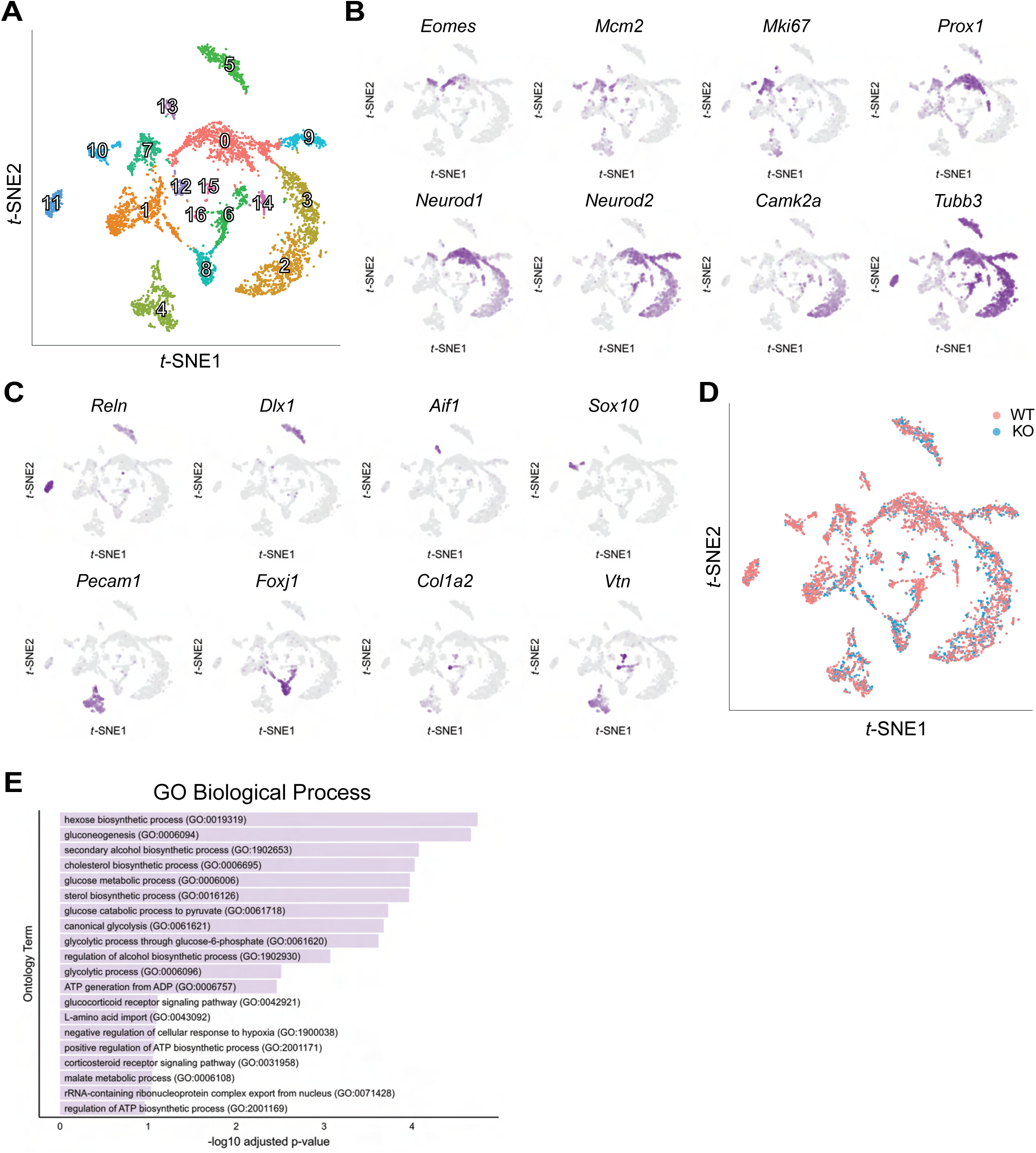
Single-cell RNA sequencing resolves the tissue heterogeneity in the developing DG. **A,** unlabeled t-SNE plot of DG cells from *hGFAP-Cre;Kdm6b^F/F^* and control mice. **B-C,** t-SNE plot of well-known marker expression for each cell type. **D,** t-SNE plot of DG cells based on the genotype of mice. **E,** Gene ontology terms identified for statistically significant down-regulated genes in *Kdm6b*-deleted NSCs.

